# Nine high-quality *Anas* genomes provide new insights into *Anas* evolution and domestication

**DOI:** 10.1101/2024.03.17.585367

**Authors:** Zhou Zhang, Zijia Ni, Te Li, Mengfei Ning, Chuze Gao, Jiaxiang Hu, Mengying Han, Jiawen Yang, Fusheng Wu, Li Chen, Lizhi Lu, Zhongzi Wu, Huashui Ai, Yinhua Huang

## Abstract

The evolutionary origin and genetic architecture of domestic animals are becoming more tractable as the availability of more domestic species genomes. Evolutionary studies of wild and domestic organisms have yielded many fascinating discoveries, while the stories behind the species diversity of *Anas* or the domestication of duck were largely unknown. Here, we assembled eight chromosome-level *Anas* genomes. Together with our recently available Pekin duck genome, we investigated *Anas* phylogeny, genetic differentiation, and gene flow. Extensive phylogenetic inconsistencies were observed in *Anas* genomes, particularly two phylogeny conflicts between autosome and Z-chromosome. However, the Z chromosome was less impacted by introgression and more suitable to elucidate phylogenetic relationships than autosomes. From the Z-chromosome perspective, we found that the speciation of *Anas platyrhynchos* and *Anas zonorhyncha* accompanied with female-biased gene flow, and remodeled duck domestication history. Moreover, we constructed an *Anas* pan-genome and identified several differentiated SVs between domestic and wild ducks. These SVs might act as repressors/enhancers to regulate their neighboring genes (i.e., *GHR* and *FER*), which represented the promising “domestication genes”. Additionally, *Anas* genomes were found being presented LTR retrotransposon bursts, which might largely contribute to functional shifts of genes involved in duck domestication (i.e., *MITF* and *IGF2BP1*). This study opens a new window to unravel avian speciation and domestication from Z chromosome.

## Introduction

Understanding the evolutionary processes of organisms in natural and domestic environments is of great interest. Domesticated organisms serve as models for studying evolution (i.e., natural selection and gene flow) and rules of inheritance (1). They also provide plenty of food to human beings and contribute to the development of permanent human societies. However, domesticated organisms demonstrate limited genetic diversity due to intensive selective breeding, which hinders their trait improvements and reduces their adaptability and resilience (2–4). Recent studies have aimed to increase genetic diversity in domesticated organisms by introducing genetic resources from related wild species and exploring functional variations within those species (5–7). This approach has been challenging in animals, mainly due to the difficulties of cross-species hybridization (8), the limited species diversity of their wild relatives(9) and few functional variations being explored.

Duck, a general name for *Anas* and other Anatidae species, is globally distributed and exhibit diverse phenotypes, with the majority being migratory and sexually dimorphic. *Anas* is one of the largest avian genera (10), and domestic duck contains a large variety of breeds (https://www.fao.org/dad-is/). Duck, especially the mallard, has the extensive cross-species hybridization capabilities, i.e., 82 hybrid combinations related to mallard, and 576 hybrid combinations related to *Anas* were recorded (http://www.bird-hybrids.com/). This enables duck to have the feasibility of cross-species breeding. Therefore, duck provides a good model for unravelling the complex processes of animal phylogeny, evolution, and domestication. The mallard was suggested as the ancestor of domestic ducks, and three genes (*MITF*, *IGF2BP1* and *NR2F2*) were inferred to contribute to the formation of the Pekin duck (11–13). However, the *Anas* evolution and domestication are still unclear. Moreover, why a few wild *Anas* species (i.e., only mallard out of sympatric duck species) has been domesticated remains unknown (14).

In this study, we assembled high-quality genomes of one domestic duck and seven divergent wild ducks. Together with the recently available high-quality reference genome of Pekin duck (SKLA2.0), we performed comprehensive evolutionary analyses to understand the evolution and domestication of *Anas* species. We further constructed an SV-based pan-genome using these nine genomes and leveraged it to explore differential SVs between ducks, especially SVs diverging between domestic and wild ducks, and to investigate effect of SVs on functional genes during duck domestication. Moreover, we investigated the dynamics of transposon elements in duck genomes, with a special emphasis on their roles in duck domestication.

## Results

### High-quality genome assemblies and annotations

Genomic sequencing was performed on eight divergent ducks from the *Anas* genus, including one domestic laying-type duck *Anas platyrhynchos* (Shaoxing, SX), and seven wild ducks, namely Mallard (*Anas platyrhynchos*, MA), Chinese spot-billed duck (*Anas zonorhyncha*, SB), Pintail (*Anas acuta*, AAC), Eurasian green-winged teal (*Anas crecca*, ACR), Falcated teal (*Anas falcata*, AFA), Northern shoveler (*Anas clypeata*, ACL), and Baikal teal (*Anas formosa*, AFO) (Fig. 1*A*, *SI Appendix,* Fig. S1). The genomes were assembled using normal/ultra-long Nanopore or Pacbio HiFi reads, as well as next generation sequencing (NGS) reads, BioNano and Hi-C data (*SI Appendix*, Fig. S2*A*, *SI Appendix*, Table S1). This effort constructed eight highly continuous genomes anchored on 40 to 41 chromosomes. The contig N50 ranged from 11.9 to 32.8 Mb, and the scaffold N50 ranged from 67.2 to 77.6 Mb (Fig. 1*B*, *SI,* Table S3). Quality evaluation indicated that eight duck genomes had high BUSCO (Benchmarking Universal Single-Copy Orthologs) values (95.5-96.3%), high consensus accuracy (QV) scores (46.5-50.5) and high mapping rates (99.56-99.81%) of population NGS reads (*SI Appendix*, Fig. S3*A* and *SI,* Table S4). These observations suggested that our eight duck assemblies were of high quality, with comparable contiguity and completeness to one of the one of the highest quality avian genomes (the chicken GRCg7b).

**Figure 1.**
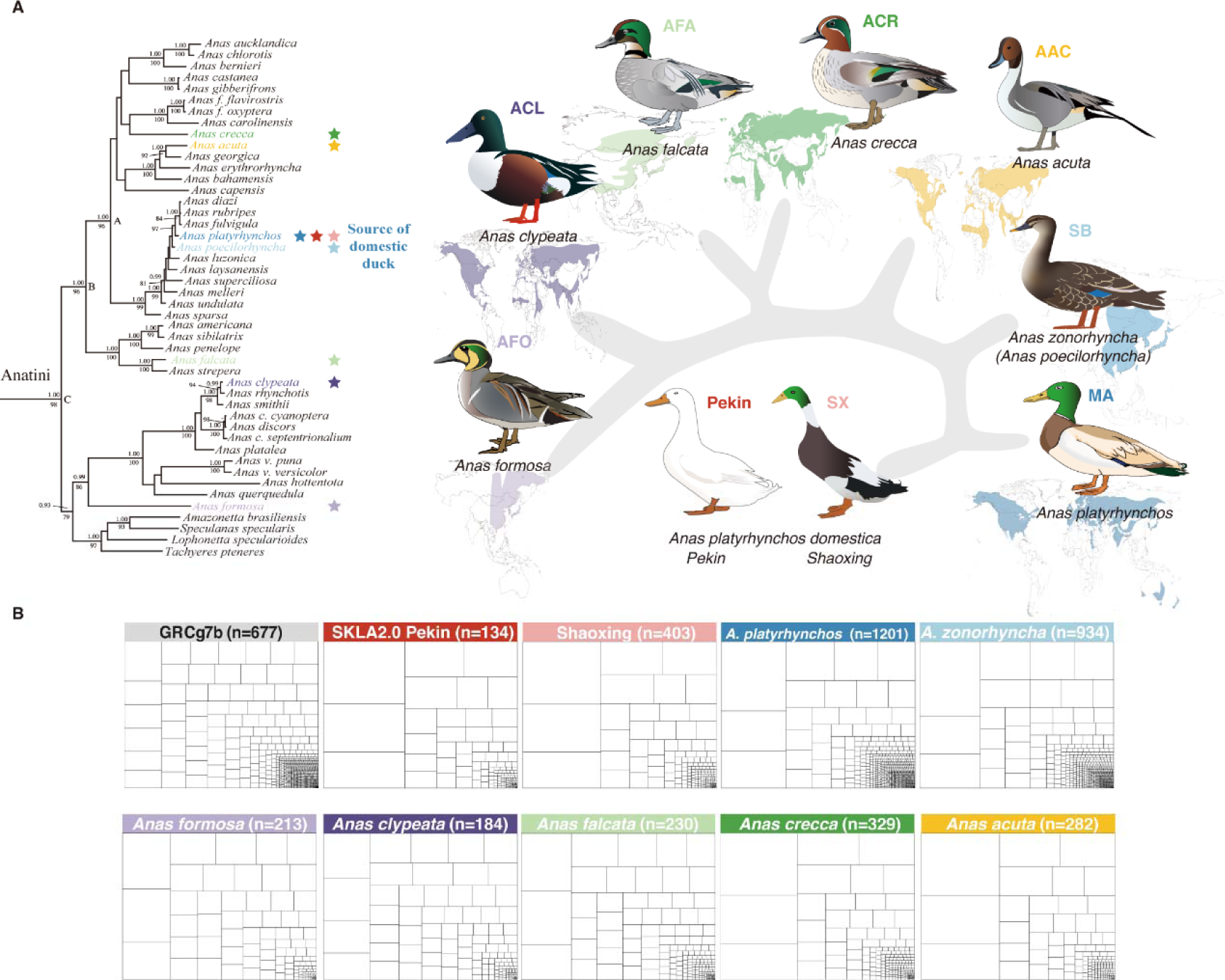
Species information and genome assembly treemaps. (A) Geographic distributions of nine (two domestic and seven wild) duck species and their male plumage during the breeding season. Source data for geographic distributions were collected from https://www.iucnredlist.org/. The Anatini phylogenetic tree in the left side and grey graph in the right side represented the phylogenetic relationships of nine ducks, which were collected from Gonzalez (Gonzalez et al. 2009) (full phylogeny available in *SI Appendix*, Fig. S1). *Anas zonorhyncha* was formerly considered a subspecies of *Anas poecilorhyncha.* (B) Contig treemaps of eight newly duck assemblies, the GRCg7b chicken genome and SKLA2.0 Pekin duck genome.

Gene annotation yielded 17,316-17,947 coding genes and 82,383-101,261 transcripts, with less than 50% overlap with repeat sequences in eight assemblies. Less than 0.5% of genes contain gaps in their 10kb flanking regions, and all transcripts have high consistency to those of Pekin duck (SKLA2.0) and GRCg7b, with at least of 99% BUSCO values (Fig. S3 *B-E*). Moreover, eight assemblies contained 13.61-20.90% repeat elements. Among them, short interspersed nuclear elements (SINEs, accounting for ∼0.05% of genome) were constant, while other four type transposable elements (TEs) were various (11.36-18.40%) across all genomes. Further analysis revealed miniature inverted-repeat transposable elements (MITEs) were substantially various, ranging from 0.09% in ACL to 6.30% in SB, and contributing to 81 Mb of sequence variation (*SI Appendix*, Table S5).

### Conservation and differentiation of *Anas* genomes

Whole genome alignment revealed *Anas* genomes were high syntenic, with a limited number of chromosome rearrangements (Fig. 2*A* and *SI Appendix*, Fig. S4) and minimal chromosome length variation (*SI Appendix*, Table S6). Subsequently, we identified highly conserved elements (HCEs) and accelerated evolution elements (ACCs) of nine *Anas* species or two domestic ducks (Pekin and SX). In total, 432 (∼0.7 Mb) domestic HCEs were observed only in domestic duck (Type I), and another 22,534 (∼35.9 Mb) domestic HCEs had orthologous sequences in at least three wild species (Type II); 2,634 (∼3.7 Mb) *Anas* HCEs were observed only in *Anas* (Type I) and another 214,201 (∼77.0 Mb) *Anas* HCEs had orthologous sequences in at least three non-*Anas* species (Type II) (Fig. 2*B*). The Type I domestic HCEs were enriched in the 0-10 kb flanking regions of genes related to cell morphogenesis and axon guidance, which are known to be associated with domestication (15), while the Type I *Anas* HCEs were enriched in the 0-10 kb flanking regions of genes involved in the Asparagine N-linked glycosylation and protein phosphorylation pathway (*SI Appendix*, Fig. S5).

**Figure 2.**
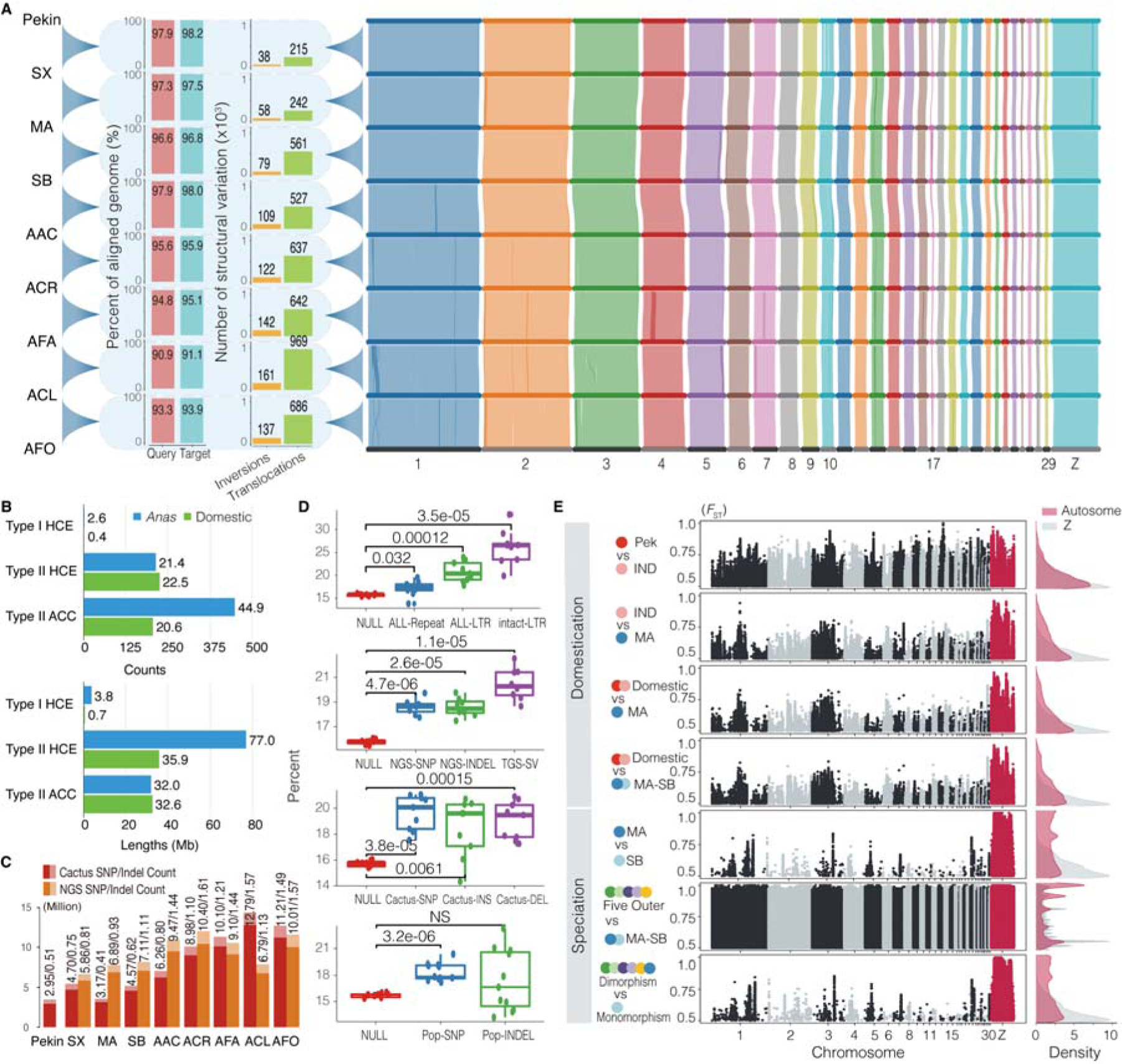
Synteny, variation, and divergence of nine *Anas* genomes. (A) The left side shows percentage coverage and number of structural variations between genomes, and the right side displays gene synteny of nine duck genomes from chr1 to chr29 and Z chromosome. Each alignment depicts the upper as the “Target” and the lower as the “Query”. (B) Statistics on HCEs (highly conserved elements) and ACCs (accelerated evolution elements) in both *Anas* and domestic ducks. (C) Comparison of variation detected with NGS data (NGS) and whole genome alignment (Cactus) in nine duck individuals. (D) Comparison in percentages of different variations in the 0-10 kb gene flanking region to percentages of the 0-10 kb gene flanking region in nine *Anas* genomes. Significance levels were calculated using a paired two-sided t-test (n=9). (E) Landscape of genomic divergence (*F*_ST_) in seven species pairs. The Five Outers are AAC, ACR, AFA, ACL, and AFO; Monomorphism refers SB, while dimorphism includes MA, AAC, ACR, AFA, ACL, and AFO.

Aligning NGS data of eight ducks to their corresponding assemblies, we identified 4.89-10.24 million SNPs and 0.78-1.54 million INDELs (referred as NGS-variants). We further extracted 2.95-12.79 million SNPs and 0.41-1.56 million INDELs (Cactus-variants) from the Cactus multi-alignment file of nine duck genomes. Many of these variants were functional mutations, such as 227-792 stop-gain and 540-1588 stop-loss in the Cactus variants (Fig. 2*C* and *SI Appendix*, Fig. S6 and S7). Notably, these variants, similar to repeat sequences, were enriched in the 0-10 kb gene flanking regions (Fig. 2*D* and *SI Appendix*, Fig. S8 *A and B*). To investigate the divergence between duck populations, the fixation index (*F*_ST_) was calculated for six population pairs. The *F*_ST_ landscapes showed that two divergent patterns were presented in *Anas*. One was only observed in the MA vs. SB pair (MA and SB primarily differed in male breeding coloration) and the dimorphism vs. monomorphism pair, where autosomes had small, but Z chromosome had large number of SNPs with high *F*_ST_ values (*F*_ST_ > 0.5). Another was presented in the remained four *Anas* population pairs, where both autosomes and Z chromosomes had many SNPs with high *F*_ST_ values (*F*_ST_ > 0.5) (Fig. 2*E*). These *Anas* divergent patterns were similar to patterns reviewed by Nosil et al (16). The former provided an example where speciation might occur with minimal genetic changes, similar to the findings in butterflies (17). While the latter suggested that both domestication and speciation, accompanied by morphological changes, occurred at numerous genetic loci, as observed in African cichlid fish (18). All comparisons, except the Five_Outer vs. MA-SB, had a relatively small number of SNPs (92 to 10,923) were nearly fixed (*F*_ST_: 0.9-1.0) (*SI Appendix*, Fig. S8*C*). This was consistent with models of polygenic adaptation from standing genetic variation (19). For ancient SNPs shared among populations, there was an increasing trend of the proportion of SNPs in the 0-10 kb gene flanking regions over increasing *F*_ST_ values (*SI Appendix*, Fig. S8 *D* and *E*). Taken together, these findings suggested that the gene regulatory regions might play a crucial role in duck speciation and domestication.

### Extensive phylogeny incongruence in *Anas* genomes

To understand the Anatidae phylogeny, we constructed a phylogenetic tree for nine *Anas* species and sixteen non-*Anas* Anatidae species with four-fold degenerate sites (4d-sites) of their genomes and estimated their divergent time using the MCMTREE (*SI,* Table S7). This analysis revealed that *Anas* species diverged from Anserinae ∼18.00 million years ago (Mya) and underwent further divergence ∼3.86 Mya, followed by the ACL and AFO divergence ∼3.31 Mya (Fig. 3*A*). It was consistent with the divergence time estimated by the PSMC analysis, which indicated that *Anas* species diverged ∼4 Mya and experienced two population declines that overlapped with the Xixiabangma Glaciation (XG, 0.8-1.17 Mya) and the Last Glacial Period (LGP, 11.7-115 kya) (Fig. 3*B*). Since the Z chromosome always plays a disproportionately role in the evolutionary process (20), we further constructed an Anatidae phylogenetic tree using 4d-sites of the Z chromosome. In general, the phylogenetic tree based on 4d-sites of the Z chromosome was consistent to those of the whole-genomes and autosomes, except for the phylogenetic position of Anserinae and the relationship between ACR and AFA (Fig. 3*C*).

**Figure 3.**
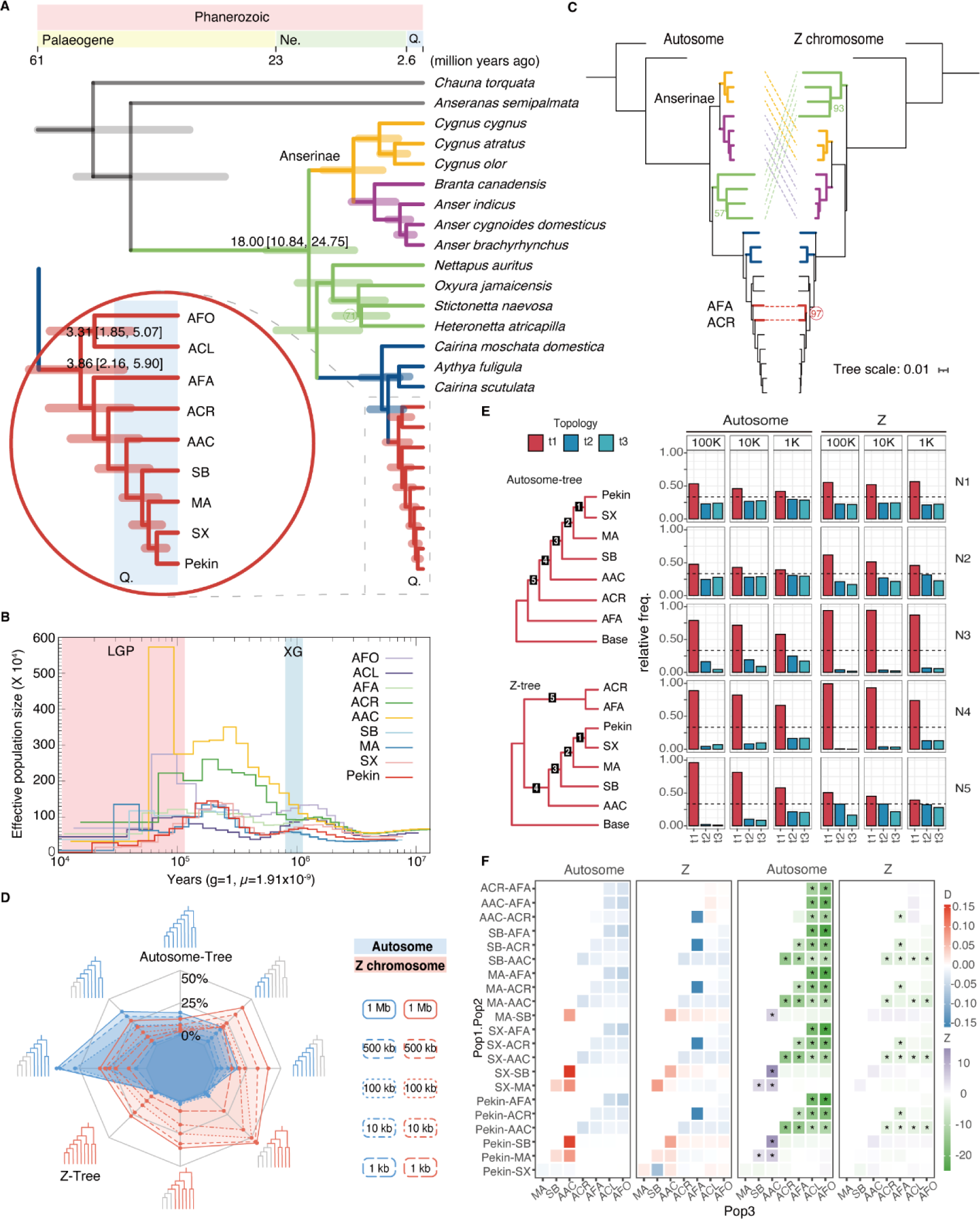
*Anas* phylogeny and extensive phylogenetic incongruence. (A) The Anatidae tree comprising nine *Anas* and 16 non-*Anas* species. The 95% highest posterior density (HPD) interval is shown after the divergence time. Bootstrap values below 100 are indicated next to the respective node. (B) PSMC estimation of nine duck genomes, with the pink shade denoting the last glacial period (LGP) and cyan shade denoting the Xixiabangma glacial (XG). (C) Two autosome-Z conflicts: (1) The relative position of the non-*Anas* duck lineage (green line) and Anserinae lineage (yellow and purple line); (2) The relationship between ACR and AFA (red line). Bootstrap values below 100 are indicated next to the respective node. (D) Comparison of autosome and Z chromosome gene trees under different window sizes (1,000, 500, 100, 10, and 1 kb). The value in each corner corresponds to the adjacent topology. The gray part of the topology indicates that it was excluded during the statistical analysis. (E) Supports for three topologies on five nodes of autosome and Z chromosome tree by 100, 10, and 1 kb gene trees. (F) *D*-stat of *Anas* genomes for gene flow detection, using *Anser brachyrhynchus* as the outgroup (pop4). The grids filled with asterisks (*) indicate that Z is greater than or smaller than 3.

The avian genome has been largely affected by incomplete lineage sorting (ILS) (21), and the *Anas* have frequently hybridized (22). Both ILS and hybridization would disrupt local phylogenetic relationships (i.e., the inconsistency of autosome and Z chromosome 4d-site trees). To assess effects of ILS on *Anas* genomes, we generated local phylogenetic trees using various genome window sizes from 1 Mb to 1 kb. As the window size decreased, the support for autosomal and Z chromosome 4d-site trees decreased from 17.73% to 0.94% and from 26.74% to 5.41%, respectively. Notably, the local phylogenetic unstableness was mainly from the “Mallard complex” (including Pekin, SX, MA, and SB) (Fig. 3*D*). However, both the coalescent trees based on window and CDS trees of the autosomes were consistent to the autosome 4d-site tree. Similarly, the coalescent trees of the Z chromosome were consistent to the Z chromosome 4d-site tree (*SI Appendix*, Fig. S9). We further investigated the phylogenetic discordance at each node of the autosome/Z coalescent trees and found most nodes had one primary (t1: 0.19-0.99) and two substantial (t2 and t3: 0.00-0.62) alternative topologies. The discordances on N1 and N4 nodes showed symmetric distributions, suggesting that ILS occurred in these two nodes. While N2 and N3 nodes displayed relatively skewed distributions, suggesting that a combination of ILS and hybridization presented in these two nodes (Fig. 3*E*). We then performed the *D*-statistic analysis on 154 four-population combinations (*SI,* Table S8). Significant signals of gene flow (|Z| > 3) were detected in 63.6% (98/154) of autosomal combinations and 28.6% (44/154) of Z chromosome combinations, including SB-domestic, AAC-domestic, AFA-AAC, ACR-AAC, and AAC-ACL (Fig. 3*F*). Compared to the autosomes, the Z chromosome had unusually higher |D| values of the combinations with ACR as pop2 and AFA as pop3. Considering the population expansion observed in the demographic history of ACR (Fig. 3*B*), and the autosomal/Z phylogenetic incongruence between ACR and AFA (Fig. 3*C*), it was likely that the ACR represented a sister species of the AFA as the Z-chromosome 4d-site tree indicated, and gene flow from an unknown species disrupted the autosomal phylogeny of ACR and caused its population expansion. In summary, hybridization and ILS resulted in extensive phylogenetic incongruences in *Anas,* making it a valuable model for phylogenetic studies.

### Speciation of the MA and SB, and inference of duck domestication history

The Z chromosome sequences showed significantly lower |Z| values and a higher proportion of the primary topology in the *Anas*, implying that the Z chromosome was less influenced by gene flow or ILS (Fig. 3 *E* and *F* and *SI Appendix*, Fig. S10*A*). We then compared the population-level phylogenetic trees constructed using SNPs from the autosomes and the Z chromosome. The neighbor-joining (NJ) trees showed that some “misplaced MAs” diverged from the other MAs (“correct MAs”) and clustered together with SBs when using autosomal SNPs, whereas all MAs were in a group and clearly diverged from SBs when using Z-chromosome SNPs (Fig. 4*A*, *SI Appendix*, Fig. S10*B*). Similarly, in the ADMIXTURE analysis, “misplaced MAs” could not distinguish from SBs when using autosomal SNPs. While all MAs clearly distinguished from SBs when using Z-chromosome SNPs, but “misplaced MAs” contained ∼15% SB ancestry, when using Z-chromosome SNPs (K=5, 6). However, neither the autosomes nor the Z chromosome of SB contained the component of MA ancestry, indicating asymmetrical introgression from SB to MA (Fig. 4*B*, *SI Appendix*, Fig. S10 *C* and *D*).

**Figure 4.**
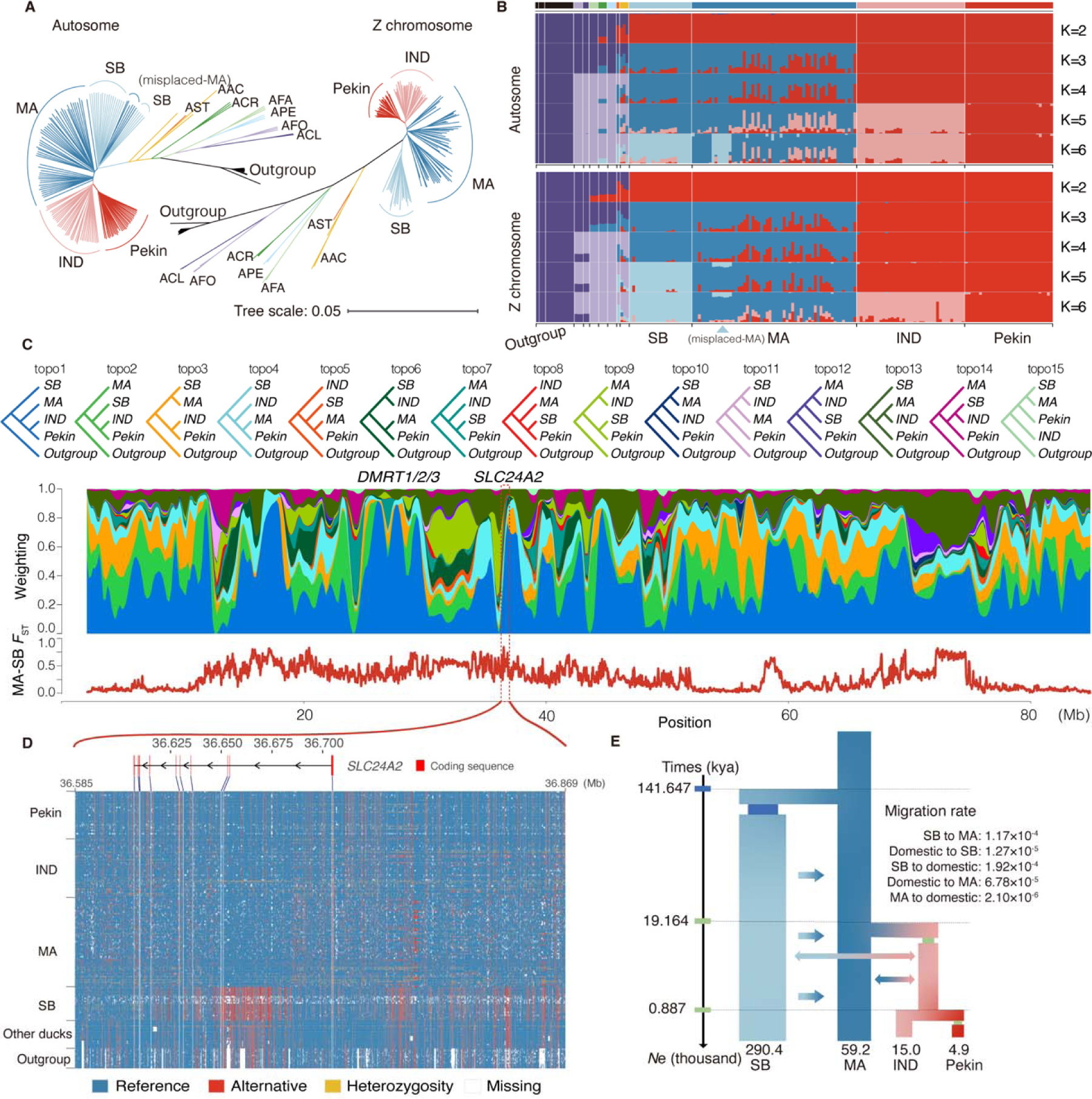
Autosome-Z chromosome comparison and domestication model. (A) Neighbor-joining tree of 180 samples from 22 duck species/breeds, goose and swan based on autosome and Z chromosome data. Different colors represent different duck species/breeds, while geese and swans are indicated with black lines as outgroups. IND: indigenous ducks, AST: *Anas strepera*, APE: *Anas penelope.* (B) ADMIXTURE results using autosome and Z chromosome data, with K=2-7. The color-coded grids at the top correspond to different groups as depicted in (A). (C) Topology weighting over the Z chromosome using 5,000-site sliding windows. The “O” are other *Anas* species except Pek, IND, MA, and SB. The different colored topologies in the bar plot correspond to the fifteen topologies indicated at the top. Genes overlapping with low-introgression regions are marked at the top of the bar plot. The bottom plot displays weighted *F*_ST_ values between MA and SB over the Z chromosome at 50 kb sliding windows. (D) SNP genotypes of the *SLC24A2* gene region in the 180 samples. (E) Best model estimated demographic scenario for duck domestication. Three time points, from top to bottom, represent MA-SB splitting, domestication, and Pekin formation events.

To further understand the resistance of the Z chromosome to introgression, we performed the topology weighting analysis. This found that 74 of 258 windows (28.7%) were resistant to introgression using the threshold of weights > 0.5 for the topology of “((((IND-indigenous duck, Pekin), MA), SB), Outgroup)”. A significant resistant region spanned ∼7.2 Mb and contained the critical *DMRT* loci known for their pivotal role in sex determination (23). Additionally, we found another resistant region that spanned ∼0.3 Mb and contained *SLC24A2* gene associated to coat and skin color (24, 25), exhibited the highest MA-SB *F*_ST_ values (Fig. 4*C*, *SI,* Table S9). However, the highly differential sites between MA and SB did not change any amino acids in *SLC24A2* (Fig. 4*D*). Further comparison suggested that “correct MAs” and “misplaced MAs” shared the same introgression pattern, suggesting that the evolutionary displacement of “misplaced MAs” was likely due to unequal ancient introgression among MA populations rather than recent introgression (*SI Appendix*, Fig. S11). Based on the above analysis, we had a clear understanding of the duck domestication history and constructed an ideal model based on: (1) the 4d-sites tree; (2) continuous asymmetric introgression from SB to MA after their split; and (3) an initial domestication stage with gene flow between MA and domestic ducks, as well as gene flow between SB and domestic ducks (*SI Appendix*, Fig. S12*A*). In the best-estimated model A, SB separated from MA ∼142 kya with continuous migration from SB to MA. Ancestors of domestic ducks diverged from MA ∼19 kya and the Pekin duck separated from the IND ∼887 years ago (Fig. 4*E*).

### Construction of *Anas* pan-genome and identification of differential SV

To create a representative reference genome, we used the Pekin (SKLA2.0) genome as the reference and constructed an *Anas* pan-genome with our new eight *Anas* genomes using the minigraph (26). This generated a pan-genome containing 698,568 nodes with a cumulative length of ∼1,263 Mb. Among them, 179,439 (25.69%) nodes with a length of ∼1,031 Mb (81.62%) were shared by nine ducks, and 259,506 (37.15%) nodes with a length of ∼124 Mb (9.82%) were sourced from eight non-reference duck genomes (*SI Appendix*, Fig. S13 *A-D*). We evaluated this pan-genome quality using HiFi and Nanopore reads of nine *Anas* individuals for genomic assembly, which showed that mapping rates were ∼98 % and ∼70 %, respectively (*SI Appendix*, Fig. S13*E*). The SVs detected by aligning NGS data of nine *Anas* individuals for genomic assembly to the pan-genome (280,430 SVs, test set) were compared with those detected by whole-genome alignment between the reference genome (Pekin duck SKLA2.0) and new eight *Anas* non-reference genomes (208,148 SVs, base set), with an F1 score of 71.3%, recall of 79.7%, and precision of 64.5% (*SI Appendix*, Fig. S13*F*). These results suggested that the pan-genome was of good quality, and suitable to identify SVs using NGS data.

To investigate how SVs contribute speciation and domestication, we identified SVs by aligning NGS reads from 200 individuals representing 22 duck species/breeds to the above *Anas* pan-genome. The overall genotyping rate for SVs in 234,931 loci was 0.70 across all 200 samples. Among these SVs, 22,072 were shared by all *Anas* species, while 5,261, 12,156, 14,764, 21,170 and 20,612 were specifically present in the ACA, ACR, AFA, ACL, and AFO, respectively (Fig. 5*A*, *SI Appendix*, Fig. S14). Moreover, we identified 19,976 non-domestic SVs in the MA and 15,588 non-domestic SVs in the SB, which were not present in domestic ducks (including Pekin and IND). These SVs might serve as valuable resources for increasing genetic diversity and/or improving the economic traits of domestic ducks through hybridization.

**Figure 5.**
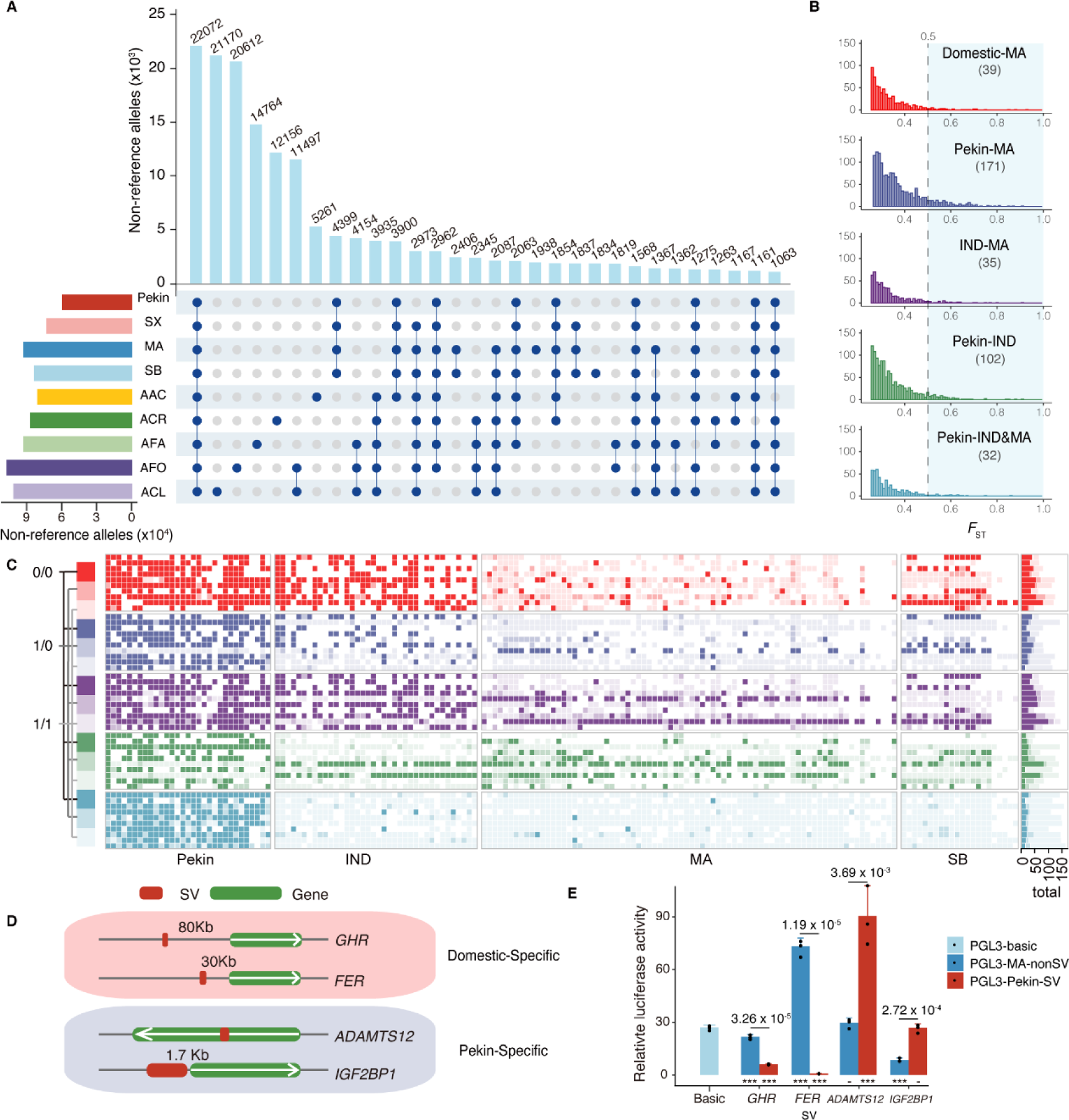
SVs distribution in ducks and their effect on gene expression. (A) Counts and overlaps of SVs among Pekin, IND, MA, and SB populations. (B) Distribution of *F*_ST_ values for SV differentiation among five population groups. (C) Genotype of SVs with the top10 *F*_ST_ values in five comparison pairs. (D) Relative positions of four highly divergent SVs and their nearby genes. (E) The dual-luciferase reporter assay of four highly divergent SVs in (D) in DF-1 cells. Data are presented as mean ± s.d. of Pekin duck allele and MA allele at these four SV loci. The *p* values above each bar plot indicate significant differences between Pekin duck allele (SV) and MA allele (Non-SV). The asterisks at the bottom represent significant differences between vector with Pekin duck allele and MA allele (two-sided t-test, three replicates for each group, *** *p* value *< 0.001*).

We further identified divergent SVs between populations by calculating *F*_ST_ values for five comparison pairs: domestic-MA, Pekin-MA, IND-MA, Pekin-IND, and Pekin-IND & MA. Each pair yielded 32-171 SVs with *F*_ST_ values greater than 0.5 (Fig. 5*B*), and most genes near these SVs showed significant differential expression between domestic and wild ducks (*SI,* Table S10 and *SI Appendix*, Fig. S15). Among SVs with top 10 *F*_ST_ values in each pair, one domestic-MA divergent SV was located at ∼80 kb upstream of the *GHR* (Growth Hormone Receptor) gene, a critical gene for body growth (27). Another domestic-MA divergent SV was in ∼30 kb upstream of the *FER* gene, which is associated with the retinal development (28). Two Pekin-specific SVs were identified in proximity to genes associated with feather color (*MITF*) (13) and chondrogenesis (*ADAMST12*) (29), respectively (Fig. 5 *C* and *D* and *SI Appendix*, Fig. S16 and S17). The *GHR, FER,* and *ADAMST12* genes showed significant differential expression between domestic and wild ducks (*SI Appendix*, Fig. S15 and S16), and the Dual-Luciferase Reporter assays showed that DF-1 cells (chicken fibroblast cell) expressing SV allele of Pekin duck had significantly higher or lower luciferase activity compared to DF-1 cells expressing SV allele of MA (Fig. 5*E*). These observations suggested that these differential SVs might have played an important role in the domestication of ducks by altering gene expression.

### Potential effect of LTR-RT burst on duck domestication

LTR-RT (long terminal repeat retrotransposons), an important source of SV, affects the host genome in many ways, such as regulating gene expression (30). Here, we observed two LTR-RT bursts occurred during the evolution of *Anas* species. The recent burst (∼122 kya) occurred in domestic ducks and their closely related species (Pekin, SX, MA, and SB), and the ancient burst (∼924 kya) occurred in other five distantly related species of domestic duck (Fig. 6*A*). Detailed analysis showed that each *Anas* genome had a specific set of intact LTR-RTs (intact-LTRs), ranging from 159 to 665, and the Pekin duck specifically shared 51 intact-LTRs (the domestic intact-LTRs) with SX, indicating that the recent LTR-RT burst was sustained during duck domestication (Fig. 6*B*, *SI Appendix*, Table S11). The divergence times of specific intact-LTRs were significantly shorter than those of non-specific intact-LTRs in each genome, consistent with the fact that specific LTRs appeared later (Fig. 6*C*). A phylogenetic tree of LTR sequences revealed six clades contained large number of expanded LTR-RTs and had short inner branch lengths. This was consistent to appearance of the recent burst peak of LTR-RTs (Fig. 6 *D* and *E*), implying that these clades might represent the recent burst event. Moreover, we characterized flanking sequences of intact-LTRs from Pekin ducks in four duck populations (Pekin, IND, MA, and SB). This found that these regions had an extremely low density of SNPs. Among them, the flanking regions of domestic intact-LTRs exhibited significantly higher *F*_ST_ values between MA and domestic ducks (Fig. 6 *F* and *G* and *SI Appendix*, Fig. S18).

**Figure 6.**
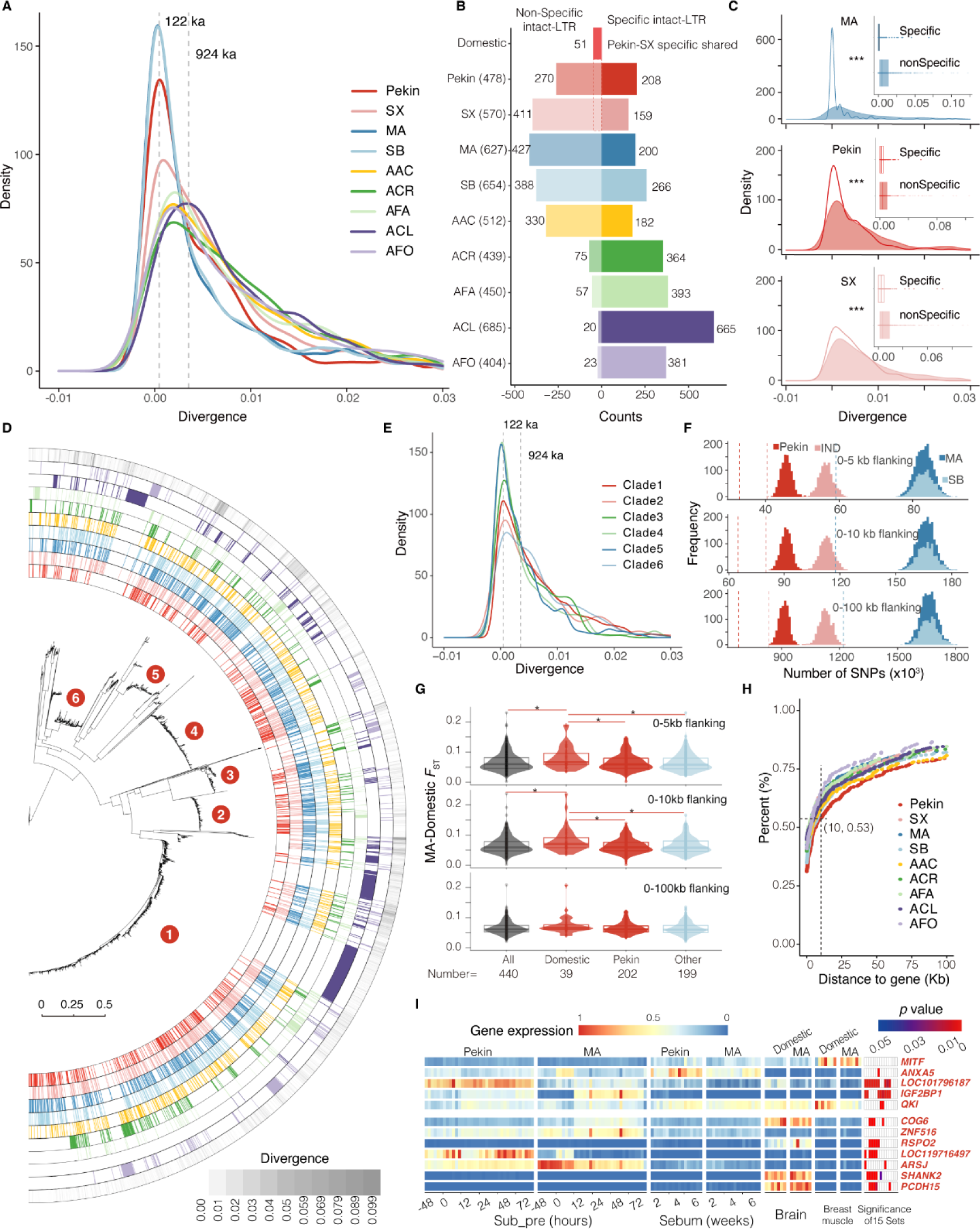
LTR-RT burst in *Anas* genomes and its effect on gene expression. (A) Distribution of LTR sequence divergence. Ages at two peaks were converted from peak values based on a mutation rate of 1.9 × 10^−9^ substitutions per site per year. (B) Category and number of intact-LTRs. (C) Distribution of LTR sequence divergence for two types of intact-LTRs; *** indicates *p* value *<* 0.001. (D) Phylogenetic tree based on two-sided LTR sequences of intact-LTRs. The outermost circle represents the degree of LTR sequence divergence, and the inwards represent the LTR positions of the different species. (E) Distribution of LTR sequence divergence from six clades in (D). (F) Comparison of actual SNP frequency (dash line) in the flanking regions (0-5, 0-10, and 0-100 kb) of intact-LTRs and the distribution of SNPs frequency (histogram) when the flanking regions of intact-LTRs in four duck specie were randomly permuted among genomes for 1,000 times. (G) Mean SNP divergence (*F*_ST_) at the flanking regions (0-5, 0-10, and 0-100 kb) of each LTR from four types of Pekin intact-LTRs; * indicates *p* value *<*0.05. (H) Cumulative plot showing the percentage of intact-LTRs to gene distance. (I) Normalized FPKM values and *p* values for differential expression of genes of interest.

Next, we investigated effect of LTR bursts on gene function. It found that more than 53.1% of the intact-LTRs were in genes or the 0-10kb gene flanking regions (Fig. 6*H*). In Pekin duck genome, 208 Pekin-specific intact-LTRs were close to 153 genes and 51 domestic intact-LTRs were close to 31 genes (*SI,* Table S12). Among these genes, 67.9% (125/184) showed differential expression between domestic and wild ducks (*SI Appendix*, Fig. S19*A*). We then manually selected four Pekin-specific intact-LTRs and seven domestic intact-LTRs for further analysis. One Pekin-specific intact-LTR resulted in the Pekin-specific SV were in the intron of the *MITF*. While another resulted in an SV located in the 13.1 kb upstream of the gastric inhibitory polypeptide (*LOC101796187*) gene, which might stimulate insulin secretion (31), and in the 1.7 kb upstream of the well-known body-weight related gene (*IGF2BP1*) (13, 32) (*SI Appendix*, Fig. S19*B* and S20). Further analysis indicated that this SV could significantly increase the luciferase activity in DF-1 cells (Fig. 5*E*). It was consistent to that Pekin ducks had a higher expression of *IGF2BP1* than MAs (Fig. 6*I* and *SI Appendix*, Fig. S19*C*), supporting that this SV might be a causative variant for body weight. The remaining two Pekin-specific intact-LTRs were close to genes involved in egg production (*ANXA5*) (33) and muscle development (*QKI*) (34), respectively. Seven domestic-specific intact-LTRs were close to genes associated with muscle development (*SLC24A3*), adipogenesis (*RSPO2*), and the hearing system (*PCDH15*) (35–37). In summary, these findings highlighted important roles of LTR-RTs in the evolutionary trajectory and domestication of ducks, through generating SVs and regulating gene expression. It might, in return, enhance ducks’ adaptability to domestic environments.

## Discussion

The evolutionary process is quite complicated, affecting by multiple factors such as gene flow, selection and ILS (38). Here, we generated eight high-quality *Anas* genomes using advanced sequencing technologies, and revealed the extensive phylogenetic inconsistencies in *Anas* species, specifically highlighting phylogenetic conflicts between autosomes and Z chromosome (Fig. 3*C* and Fig. 4*A*). By comparing the evolutionary characteristics (i.e. phylogeny, ancestral components) of the Z chromosome to those of autosomes, we inferred the hybrid speciation of ACR, where the hybrid speciation is increasingly recognized as a creative evolutionary force contributing to species adaptation and speciation (39). We also proposed that the MA-SB speciation was a sympatric speciation accompanied by sexual selection-mediated female-biased gene flow (*SI Note 1*). The phylogenetic inconsistencies between the autosomes and the Z chromosome might be partially attributed to the Z chromosome’s resistance to introgression (Fig. 4*C*). This suggested that, for species with frequent cross-species hybridization, such as birds (40), the Z chromosome with less introgression was more suitable for elucidating phylogenetic relationships and the processes of domestication than the autosomes. Recent avian phylogeny were estimated based on resequencing data or genomes with relatively low-quality and/or lack of the Z chromosome sequence (41, 42). Such analysis might lead to biased or controversial avian phylogeny (43). The future ability to generate high-quality avian genomes including the Z chromosome sequence and to compare the evolutionary history of the autosomes and Z chromosome will certainly extend our knowledge of avian evolution.

Spatiotemporal patterns of domestication, domestication genes and factors behind a few wild species being domesticated are three key questions in the area of domestication studies (1, 44). In this study, we have performed meaningful exploration to partially answer these three questions in *Anas* species. Firstly, we found that multiple gene flows, particular the introgression from SB to domestic duck, were occurred in duck domestication, and further remodeled a clearer duck domestication history when compared to previous studies (12). We obtained the best-estimated model (model A, with lowest AIC values) of duck domestication, assuming that ancestor of domestic ducks diverged from their descended MA ∼19 kya. This was consistent with the identified bottleneck event (*SI Appendix*, Fig. S21). However, it is important to acknowledge that the possibility of alternative scenarios (model B, J and K) cannot be entirely excluded (*SI Appendix*, Fig. S12 *B* and *C*). More evidence, especially fossil records, is required to confirm the inferred timeline of duck domestication. Secondly, we identified several SVs that appeared in nearly all domestic ducks but were rarely observed in wild ducks. Two of these SVs showed regulatory effects on their downstream genes in vitro. These observations together with neighbor genes (*GHR*, *FER*) of these two SVs showed significantly differential expression between domestic and wild ducks, suggests that these regions might have been under strongly selected and contributed to phenotypic changes (such as body growth) in domestic ducks. Thirdly, our study highlighted an LTR-RT burst in the wild ancestors of domestic ducks, and the evolutionary signatures, such as SNP density, around intact-LTR regions. Previous studies have shown that LTR-RT burst was related to the environment adaption in birds (45), and here we found that LTR-RT burst might be involved in duck domestication. For example, some Pekin or domestic specific intact-LTRs potentially regulate the expression of their neighbor genes associated with animal domestication, including *IGF2BP1* for increasing body size (13), *MITF* for feather color changing (13), *PCDH15* for adapting environment (35) and *ANXA5* for enhancing productive ability (33). These insights might help to explain why MA, compared to other wild duck species, has been successfully domesticated by humans. Of course, further experimental evidence is required to unravel how these intact-LTRs regulated “domestication genes” and contributed to duck domestication.

Of special interest to agriculture and medicine is the fact that ducks hold significant importance as a domestic animal and one of the principal natural reservoirs for influenza A viruses. In this study, we detected large number of cross-species genetic variants (2.95-12.79 million SNPs, 0.41-1.57 million INDELs and 0.23 million SVs) in *Anas* using our eight high-quality genomes, together with our recently available Pekin duck genome. Among them, 135,311 SVs observed in wild ducks were overlapped with 11,748 genes. In particular, these SVs overlapped with the CDS region of 317 genes, enriched in carbohydrate metabolism, regulation of anatomical structure size et al (*SI Appendix*, Fig. S22). Since domestic ducks might extensively hybridize with wild ducks (http://www.bird-hybrids.com/), such substantial genetic variants of wild ducks might be used to increase genetic diversity of domestic ducks. It would in return magnify phenotype variation and accelerate duck breeding. Moreover, our eight chromosome-level *Anas* genomes, together with their reference gene sets, provide resources for fine charactering interaction between host and influenza viruses.

## Materials and Methods

### Sample collection and data generation

One female domestic duck (SX) for genomic sequencing was collected from Zhejiang Guowei duck farm, Zhejiang, China. Seven female wild ducks for genomic sequencing were collected from two duck farms with available hunting licenses: Zhejiang Aoji Duck farm in Zhejiang, China; Longyao farm for special economic animals in Shandong, China.

Genomic DNA was isolated from fresh blood and RNA were isolated from four or five tissues of eight ducks. Normal ONT (Oxford Nanopore Technologies) long reads for six samples, ONT ultralong reads and full-length transcripts for all eight ducks were generated using the Nanopore PromethION sequencer and HiFi reads for two samples (MA and SB) were generated using the PacBio Sequel II sequencer (SI Appendix, Table S1). Optical molecules of high-quality genomic DNA (labeled restriction enzyme DLE1) were produced using the BioNano Genomics (BNG) instrument Saphyr. Short paired-end reads of the genomic DNA and the Hi-C libraries for eight duck samples were sequenced using the Illumina NovaSeq 6000 or MGISEQ-2000 platform (SI Appendix, Table S1). All library preparations and sequencing were conducted at the Genome Center of GrandOmics Bioscience Co., Ltd. (Wuhan, China). The ONT reads with quality scores ≥ 7 were utilized for subsequent analyses, and the HiFi reads were extracted from the bam files using the ccs (v3.4.1, https://github.com/PacificBiosciences/ccs) software with default parameters. Unless otherwise stated, all NGS DNA and RNA data used in subsequent analyses underwent quality control by fastp (v0.20.0) with default parameters (46).

### Genome assembly and annotation

Briefly, six ducks (SX, AAC, ACR, AFA, ACL, and AFO) without HiFi data were assembled and polished by using Nextdenovo (v.2.2) and NextPolish (v1.1.0) (47). The contigs were hybrid assembled with BioNano maps using the BioNano Solve (v3.3) (https://bionano.com/software-downloads/). Scaffolds were then assigned to chromosomes using Hi-C-based proximity-guided assembly, followed by manual adjustment using juicerbox (v1.11.08) (48) and 3d-dna (v180922) (49) software. After that, gaps in the assembly were filled using TGS-Gapcloser (v1.1.1) (50). For two duck species (MA and SB) with HiFi data, their assembly was performed using Hifiasm (v0.16.0) (51), while the other steps are same to above six ducks (*SI Appendix*, Fig. S2*A*). For genome annotation, we identified transposable elements using the EDTA (v.1.9.6) (52) and annotate repeat sequences using the RepeatMasker (v4.1.0) (https://www.repeatmasker.org/-RepeatMasker/). We then integrated information of homology prediction, de novo prediction and RNA sequences to predict protein-coding genes. In brief, RNA evidence from three sources was preprocessed. Protein sequences of human (GRCh38), mouse (GRCm39), chicken (GRCh6a) and duck (ZJU1.0) downloaded from the NCBI database, together with our recently Pekin duck reference protein, were used as homology evidence. The RNA-derived gtf files and homologous protein sequences were then input into the MAKER (v3.01.03) (53) pipeline to obtain the annotation of protein-coding genes. On the other hand, transcript models were extracted directly from long-read transcripts, and the Pekin duck reference gene set (SKLA2.0) were lift to our assemblies using the official command of Liftoff (v1.6.1) (54). Finally, the gtf files generated by MAKER, Liftoff, long-read transcripts, and repeat annotation were combined using in-house scripts (*SI Appendix*, Fig. S2*B*).

Detailed descriptions of genome assembly and annotation were provided in ***SI Note 2***.

### Evaluation of assembly and annotation

The quality of the genome assembly and annotation were assessed as the following: Firstly, eight *de novo* assemblies, together with Pekin duck (SKLA2.0) genome and chicken (GRCg7b) genome, were blast against the aves_odb10 dataset containing 8,338 conserved protein models, using BUSCO (v4.0.6) program in genome mode, with default settings. Secondly, eight ducks’ NGS data were mapped to their assemblies to identify variants using the HaplotypeCaller module of GATK (v4.1.8.0) (55). After quality control (see variants calling part), the homozygous variants were extracted and used to calculate quality value (QV) of genome assemblies. Thirdly, predicted proteins of the above assemblies were blast against the aves_odb10 dataset in protein mode to evaluate annotation quality. Finally, the predicted proteins of the above eight assemblies (Query) were compared to the proteins of the GRCg7b and SKLA2.0 annotations (Target) using BLASTP (2.9.0+) (56).

### Comparative genomic analysis

The guide species tree of nine *Anas* and sixteen non-*Anas* species (*SI,* Table S7) was generated by OrthoFinder (v2.5.1) (58), and then the whole-genome alignments (WGA) were performed using the Cactus (v1.2.0) (57) with the soft-masked genomes. The resulting hal file was converted to the maf files using hal2maf(59) program (--onlyOrthologs --noAncester --nodupes --refGenome SKLA2.0), and subsequently filtered using mafFilter program (https://anaconda.org/bioconda/ucsc-maffilter) with the -minCol=100 option. After that, the 4d-site msa (multi-sequence alignments) files were obtained from the filtered maf files according to Chen et al (60), and removed gaps using Gblock (61). Autosome and/or Z chromosome 4d-site msa files were used to construct phylogenetic trees with RAxML-ng (v1.0.1) (62) (--model GTR+G --threads 100 --all --bs-trees 1000).

Genomes were sequentially aligned to each other using minimap2 (version 2.17-r941)(63) with parameters: -ax asm5 –eqx. The genomic synteny and structural rearrangements were then identified using SyRI (v1.6) (64) and visualized using plotsr (65). For gene synteny, the longest transcripts of all species were aligned using BLASTP with parameters of e-value < 1e^-10^, -max_target_seqs 1. Synteny analysis was then performed using the MCScanX (66) package with default settings, and gene synteny was generated using an online website (https://synvisio.github.io/) (67).

HCEs analysis was carried out similar as previous study (60). In our analysis, sixteen non-*Anas* species were used as outgroups to identify the conserved elements of *Anas,* and sixteen non-*Anas* species together with seven wild duck species were used as outgroups to determine the conserved elements of domestic ducks (Pekin and SX).

### Divergence time calibration

Species divergence times were calibrated using MCMCTREE in PAML (Version 4.9d) (68). Five external calibration times from Prum (41) were added to the autosome 4d-site tree. The MCMCTREE run involved increasing the burn-in steps and sampling step size until convergence was achieved, indicated by an Effective Sample Size (ESS) greater than 200. Finally, the posterior distribution of divergence times based on 50000 samples was obtained by MCMC (Markov chain Monte Carlo) sampling, setting the burn-in steps to 2×10^7^ and sampling every 1,000 steps.

### Demographic history inferences and remodeling

To infer the *Anas* demographic history, a PSMC (69) analysis was performed according to official guidelines. A generation time (g) of one year and a mutation rate per generation of µ=1.91×10^-9^ were used (70) to draw the demographic history using psmc_plot.pl script. To infer the recent demographic history of Pekin, IND, MA, and SB, ten individuals (five males and five females) with less introgression was selected from the population VCF file, according to the admixture results. To avoid the influence of ancient historical events on the inference of recent demographics, variants detected in the other five wild ducks were filtered out using VCFtools (http://vcftools.sourceforge.net). SMC++ ( v1.15.2) (71) analysis was conducted for autosomes according to official procedures with the same generation time and mutation rate as the PSMC analysis.

To remodel duck domestication history, different domestication models were tested using fastsimcoal2 (72). First, the VCF file for SMC++ analysis was converted to a site frequency spectrum (SFS) file using EasySFS (https://github.com/isaacovercast/easySFS). After that, models with the following parameters: -n 100000 -M -c12 -q -multiSFS were evaluated under 100 run times using fastsimcoal2, and the model with the lowest Akaike information criterion (AIC) value was selected as the preferred model. Strategies and codes for domestication model analyses were referred an online website (https://speciationgenomics.github.io/).

### Introgression and incomplete lineage sorting analyses

For introgression analysis, the MAF files were transformed into Variant Call Format (.vcf) by mafFilter, and subsequently files were converted into a geno file using vcf2eigenstrat.py script from gdc-master (https://github.com/mathii-/gdc). D-statistics (so called ‘ABBA-BABA’ test) for different species combinations were carried out using qpDstat module from Admixtools (v7.0.2) (73), and topology weighting analysis was conducted with reference to the online manual (https://github.com/simonhmartin/genomics_general).

For ILS analysis and gene tree analyses, WGAs files were split into sliding windows using msa_split from PHAST (v1.4) (74). Small windows (10 kb, 1 kb) with gap ratios >10% were filtered out. The coalescent analysis were performed in accordance with Feng’s study (38). Briefly, 1,066 (1 Mb), 2,060 (500 kb), 7,298 (100 kb), 61,902 (10kb), and 584,729 (1 kb) qualified autosomal window trees, and 86 (1 Mb), 171 (500 kb), 505 (100 kb), 4,970 (10 kb), and 50,144 (1 kb) qualified Z chromosome window trees were input into ASTRAL-III (v5.6.2) (75) to obtain the coalescence species tree. The phylogenetic discordance was calculated with DiscoVista (v1.0) (76) in different window sizes, taking the 4d-site tree as the reference tree.

### Variants calling

For individuals with genome assemblies, SNPs and INDELs were extracted from the VCF file, which were generated in genome assessment (referred as NGS-SNP and NGS-INDEL). Insertions (Cactus-INS), deletions (Cactus-DEL), and SNP (Cactus-SNP) from WGAs for each species were extracted using halBranchMutations (59) program from the cactus toolkit. Long reads of genome assembled individuals were aligned to their corresponding assemblies using minimap2, and structural variants (TGS-SV) were identified using SVIM (v1.4.2) (77) with default parameters and filtered according to the recommendations of the authors. Variants were annotated using ANNOVAR (78) and visualized using maftools (R package, version 2.4.12) (79).

Population-SNP and population-INDEL were detected with clean reads of 174 available samples (13, 80–82) and 26 samples sequenced in our study (*SI,* Table S13) by aligning to the reference genome (SKLA2.0 and the W chromosome from ZJU1.0) using BWA (0.7.17-r1188) (83) with default parameters. Joint SNP calling for 180 out of the 200 samples was performed following GATK best practices workflow suggested on the official website. SNPs were filtered with “QualByDepth (QD) < 2.0, mapping quality (MQ) < 40.0, Fisher Strand (FS) > 60.0, StrandOddsRatio (SOR) > 3.0, MQRankSum < −12.5, ReadPosRankSum < −8.0”, and INDELs were filtered with “QD < 2.0, FS > 200.0, SOR > 10.0, MQRankSum < −12.5, ReadPosRankSum < −8.0”. While the SNP calling for additional 20 public MA samples from Zhang el al (80) was conducted by the GATK HaplotypeCaller module based on the variant sites from the joint calling, and with an additional filter to remove variants in regions with a mean depth of less than 5×.

### Distribution analysis of repeats and variants

The nine *Anas* genomes were divided into gene (UTR not included), CDS, intron, and gene flanking regions (0-10 kb, 10-20 kb, 20-30 kb, 30-40 kb, 40-50 kb, and >50 kb). Various genomic features (all-repeats, all-LTR, intact-LTR, HCE, NGS-SNP, NGS-INDEL, TGS-SV, Cactus-SNP, Cactus-INS, Cactus-DEL, population-SNP, and population-INDEL) were considered. The relative distance from genomic features to the gene and the distribution of genomic features were obtained using bedtools (v2.30.0) (84). Paired sample t-tests were conducted to determine whether genomic feature in specific region was disproportionately compared to the null distribution (percentages of different genome regions in the genome). A chi-square test was performed on 2 × 2 contingency tables to analyze the distributions of HCE features. The gene enrichment analysis was performed on the online website (https://metascape.org/).

### Population structure analysis

Autosomal and Z-linked SNPs were filtered by PLINK2.0 (85) (--maf 0.01 --geno 0.2 --indep-pairwise 50 5 0.5). The sex of samples were determined using the ‘plink –sex-check’ command, females’ heterozygous sites and the 0-2 Mb region (containing the pseudo-autosome) on Z chromosome were removed. Distance metrics between individuals were obtained using the ’plink –distance square 1-ibs flat-missing’ command, and subsequently used to construct autosomal and Z chromosome phylogenetic trees with the phylip (v3.697) neighbor program. Phylogenetic trees were visualized on an online website (https://itol.embl.de/). Using the same dataset, autosome and Z chromosome admixture were quantified among all samples using ADMIXTURE (v1.3) (86) for possible group numbers from 2 to 7, and the --haploid=”male:X” parameter was added in the run for Z chromosome.

### Long-terminal repeats retrotransposon analysis

Intact-LTRs on chromosomes were collected from the above EDTA results. The divergence of two-sided LTR sequences for intact-LTRs was converted to the insertion time of intact-LTR using formula: *Divergence*/(1.91×10^-9^ x2). Shared Intact-LTRs were identified by aligning Intact-LTRs to target genomes with thresholds of length > 11 kb and identity > 0.95. Specific LTR-RTs referred to LTR-RTs that existed only in one genome; specific shared LTR-RTs meant LTR-RTs shared by an exact number of genomes.

LTR sequences of intact-LTRs (*Gypsy*-type) were aligned using MAFFT (v7.475) (87) with default parameters. Phylogenetic tree of LTR sequences was generated using iqtree2 (v2.1.2) (88) with the K80 model. Permutation was performed by shuffling the genome blocks of different types of intact-LTR and calculating the SNP density or *F*_ST_ values 1,000 times. Welch’s two-sample t-test was conducted using R to compare the divergences or signatures of different intact-LTRs.

### Pan-genome construction and evaluation

An SV-based graph pan-genome was constructed by running minigraph (version 0.18-r538) to integrate eight new genomes and the SKLA2.0 Pekin genome. The graph in GFA format was converted to graph indexes using the vg (v1.43.0) autoindex command. The source of the nodes in the pan-genome was identified by referring to previous studies (89).

For pan-genome evaluation, long reads from the above nine ducks were aligned to the graph genome using the giraffe module of the vg program with the -align-from-chains option. The aligned gaf file was subjected for quality control, and the mapping rate was calculated using the R script from Liao et al (90). NGS reads of these nine ducks were aligned to the pan-genome to detect SVs using the giraffe, pack, and call modules of the vg program. The ‘PASS’ SVs from these nine ducks were merged into one SV dataset (test SV set) using BCFtools (v1.15.1) (91). On the other hand, eight duck assemblies were aligned to the reference (SKLA2.0) with minimap2, and SVs were identified using paftools (2.24-r1122). SVs with a length ≥ 50 bp were merged using jasmine (v1.1.4) (92) to obtain a SV dataset (base SV set), and further compared to the test SV set using Truvari (v3.5.0) (93) under defaults.

### SV divergence analysis

All SVs of the 200 samples were identified using the same methods as the above nine individuals with NGS data. SV *F*_ST_ was calculated for five comparison pairs (Pekin vs. IND, Pekin vs. MA, IND vs. MA, Pekin & IND vs. MA, Pekin & IND vs. MA & SB) using the following formula:

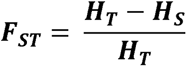

Where:

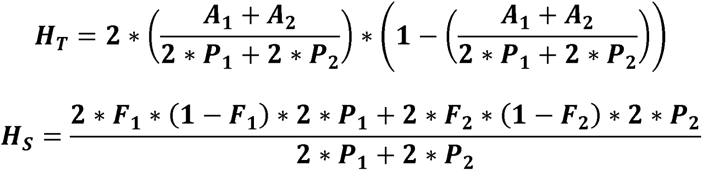

H_T_ and H_S_ are the heterozygosity of the total population and the average heterozygosity of subpopulations expected under Hardy–Weinberg equilibrium; A_1_, A_2_ are the number of SV frequencies in population1, population2; and P_1_, P_2_ are the size of the population1, population2; F_1_, F_2_ are the SV frequency in population1, population2.

### Luciferase reporter assays on highly divergent SVs

Four SVs were tested by PCR amplification using genomic DNA (*SI Appendix*, **Fig.** S23 and *SI Appendix*, **Table** S14), and their purified PCR products used to construct six pGL3-Pekin-SV vectors, six pGL3-MA-control vectors, and one pGL3-basic control vector. The dual-luciferase activity of vectors was evaluated in DF-1 cells. In brief, vectors were transfected into DF-1 cells with jetPRIME (Polyplus), and dual-luciferase activity was measured using the Dual-Glo Luciferase Assay kit (Promega) after a 48-h incubation with three biological replicates. Differences in fluorescence intensity between vectors in DF-1 cells were compared using the two-sided t-test.

### RNA-seq analysis

Available clean RNA-Seq reads of 185 samples (11, 80, 94) (*SI,* Table S15) were aligned to the SKLA2.0 Pekin duck genome using HISAT2. Number of mapped reads was calculated using featureCounts (version 2.0.3) software (95) and subsequently used to count FPKM values using the Python bioinfokit (v0.9.1) package.

Fifteen pairs of RNA-seq data were used to identify different expressional genes in MA and Pekin (or domestic) (*SI Appendix*, Table S16). Samples with low correlation (R^2^ < 0.95) compared to other samples were excluded as invalid biological replicates. The fold-change and adjusted *p* value of each gene was calculated using DESeq2 (v.1.24.0) software. Gene expression was visualized using FPKM values normalized with the maximum FPKM value of each gene in fifteen RNA data.

## Acknowledgments

We thank Prof. Lusheng Huang (Jiangxi Agricultural University), Prof. Wen Wang (Northwestern Polytechnical University), Prof. Yu Jiang (Northwest A&F University) and Prof. Guojie Zhang (Zhejiang University) for discussion and comments about avian evolutionary analysis. We acknowledge Xuetao Huang (Zhejiang Guowei duck farm), Huabin Cao (Jiangxi Agricultural University), Puyan Meng (Wildlife Rescue and Breeding Center of Jiangxi Province) for sample collections. The sequencing of the duck genomes was funded by the National Waterfowl-Industry Technology Research System (CARS-42) and the National Science Foundation of China (32172716). This work was supported by China Agriculture Research System (No. CARS-35) and the National Key Research and Development Program of China (2022YFF1000100).

## Notes

### Competing Interest Statement

The authors have declared no competing interest.

## References

1. L. Andersson, M. Purugganan, Molecular genetic variation of animals and plants under domestication. Proc. Natl. Acad. Sci. U.S.A. 119, e2122150119 (2022).

2. B. T. Moyers, P. L. Morrell, J. K. McKay, Genetic Costs of Domestication and Improvement. Journal of Heredity 109, 103–116 (2018).

3. F. Biscarini, E. L. Nicolazzi, A. Stella, P. J. Boettcher, G. Gandini, Challenges and opportunities in genetic improvement of local livestock breeds. Front Genet 6, 33 (2015).

4. S. Swarup, et al., Genetic diversity is indispensable for plant breeding to improve crops. Crop Science 61, 839–852 (2021).

5. D. Tang, et al., Genome evolution and diversity of wild and cultivated potatoes. Nature 606, 535–541 (2022).

6. L. Chen, et al., Genome sequencing reveals evidence of adaptive variation in the genus Zea. Nat Genet (2022) 10.1038/s41588-022-01184-y (October 28, 2022).

7. L. Shang, A super pan-genomic landscape of rice. Cell Research, 19 (2022).

8. T. E. Dowling, C. L. Secor, The Role of Hybridization and Introgression in the Diversification of Animals. Annu. Rev. Ecol. Syst. 28, 593–619 (1997).

9. K. H. Redford, N. Dudley, Why should we save the wild relatives of domesticated animals? Oryx 52, 397–398 (2018).

10. W. J. Bock, J. Farrand, The Number of Species and Genera of Recent Birds: A Contribution to Comparative Systematics’. 36 (1980).

11. F. Zhu, et al., Three chromosome-level duck genome assemblies provide insights into genomic variation during domestication. Nat Commun 12, 5932 (2021).

12. X. Guo, et al., Revisiting the evolutionary history of domestic and wild ducks based on genomic analyses. Zoological Research 42, 43–50 (2021).

13. Z. Zhou, et al., An intercross population study reveals genes associated with body size and plumage color in ducks. Nat Commun 9, 2648 (2018).

14. J. Diamond, Evolution, consequences and future of plant and animal domestication. Nature 418, 700–707 (2002).

15. Y. Hou, et al., Genome-wide analysis reveals molecular convergence underlying domestication in 7 bird and mammals. BMC Genomics 21, 204 (2020).

16. P. Nosil, J. L. Feder, Z. Gompert, How many genetic changes create new species? Science 371, 777–779 (2021).

17. N. B. Edelman, et al., Genomic architecture and introgression shape a butterfly radiation. Science, 7 (2019).

18. D. Brawand, et al., The genomic substrate for adaptive radiation in African cichlid fish. Nature 513, 375–381 (2014).

19. R. Barrett, D. Schluter, Adaptation from standing genetic variation. Trends in Ecology & Evolution 23, 38–44 (2008).

20. B. Charlesworth, J. A. Coyne, N. H. Barton, The Relative Rates of Evolution of Sex Chromosomes and Autosomes. The American Naturalist 130, 113–146 (1987).

21. A. Suh, L. Smeds, H. Ellegren, The Dynamics of Incomplete Lineage Sorting across the Ancient Adaptive Radiation of Neoavian Birds. PLoS Biol 13, e1002224 (2015).

22. P. Lavretsky, K. G. McCracken, J. L. Peters, Phylogenetics of a recent radiation in the mallards and allies (Aves: Anas): Inferences from a genomic transect and the multispecies coalescent. Molecular Phylogenetics and Evolution 70, 402–411 (2014).

23. C. A. Smith, et al., The avian Z-linked gene DMRT1 is required for male sex determination in the chicken. Nature 461, 267–271 (2009).

24. F. Wang, et al., A Genome-Wide Scan on Individual Typology Angle Found Variants at SLC24A2 Associated with Skin Color Variation in Chinese Populations. Journal of Investigative Dermatology 142, 1223–1227.e14 (2022).

25. D. Li, et al., Breeding history and candidate genes responsible for black skin of Xichuan black-bone chicken. BMC Genomics 21, 511 (2020).

26. H. Li, X. Feng, C. Chu, The design and construction of reference pangenome graphs with minigraph. Genome Biol 21, 265 (2020).

27. E. O. List, et al., Endocrine Parameters and Phenotypes of the Growth Hormone Receptor Gene Disrupted (GHR−/−) Mouse. Endocrine Reviews 32, 356–386 (2011).

28. A. W. B. Craig, R. Zirngibl, K. Williams, L.-A. Cole, P. A. Greer, Mice Devoid of Fer Protein-Tyrosine Kinase Activity Are Viable and Fertile but Display Reduced Cortactin Phosphorylation. Mol Cell Biol 21, 603–613 (2001).

29. X. H. Bai, D. W. Wang, Y. Luan, X. P. Yu, C. J. Liu, Regulation of chondrocyte differentiation by ADAMTS-12 metalloproteinase depends on its enzymatic activity. Cell. Mol. Life Sci. 66, 667–680 (2009).

30. G. Bourque, et al., Ten things you should know about transposable elements. Genome Biol 19, 199 (2018).

31. R. A. Pederson, C. H. McIntosh, Discovery of gastric inhibitory polypeptide and its subsequent fate: Personal reflections. J Diabetes Investig 7, 4–7 (2016).

32. K. Wang, et al., The Chicken Pan-Genome Reveals Gene Content Variation and a Promoter Region Deletion in *IGF2BP1* Affecting Body Size. Molecular Biology and Evolution 38, 5066–5081 (2021).

33. D. Wang, et al., Integrative analysis of hypothalamic transcriptome and genetic association study reveals key genes involved in the regulation of egg production in indigenous chickens. Journal of Integrative Agriculture 21, 1457–1474 (2022).

34. X. Chen, et al., The Emerging Roles of the RNA Binding Protein QKI in Cardiovascular Development and Function. Front. Cell Dev. Biol. 9, 668659 (2021).

35. L. Liu, et al., Template-independent genome editing in the Pcdh15 mouse, a model of human DFNB23 nonsyndromic deafness. Cell Reports 40, 111061 (2022).

36. A. Georges, et al., Genetic investigation of fibromuscular dysplasia identifies risk loci and shared genetics with common cardiovascular diseases. Nat Commun 12, 6031 (2021).

37. H. Dong, et al., Identification of a regulatory pathway inhibiting adipogenesis via RSPO2. Nat Metab 4, 90–105 (2022).

38. S. Feng, et al., Incomplete lineage sorting and phenotypic evolution in marsupials. Cell 185, 1646–1660.e18 (2022).

39. J. Ottenburghs, Exploring the hybrid speciation continuum in birds. Ecol Evol 8, 13027–13034 (2018).

40. J. Ottenburghs, R. C. Ydenberg, P. Van Hooft, S. E. Van Wieren, H. H. T. Prins, The Avian Hybrids Project: gathering the scientific literature on avian hybridization. Ibis 157, 892–894 (2015).

41. R. O. Prum, et al., A comprehensive phylogeny of birds (Aves) using targeted next-generation DNA sequencing. Nature 526, 569–573 (2015).

42. E. D. Jarvis, et al., Whole-genome analyses resolve early branches in the tree of life of modern birds. Science 346, 1320–1331 (2014).

43. M. P. Simmons, M. S. Springer, J. Gatesy, Gene-tree misrooting drives conflicts in phylogenomic coalescent analyses of palaeognath birds. Molecular Phylogenetics and Evolution 167, 107344 (2022).

44. G. Larson, et al., Current perspectives and the future of domestication studies. Proceedings of the National Academy of Sciences 111, 6139–6146 (2014).

45. E. Carotti, et al., LTR Retroelements and Bird Adaptation to Arid Environments. IJMS 24, 6332 (2023).

46. S. Chen, Y. Zhou, Y. Chen, J. Gu, fastp: an ultra-fast all-in-one FASTQ preprocessor. Bioinformatics 34, i884–i890 (2018).

47. J. Hu, J. Fan, Z. Sun, S. Liu, NextPolish: a fast and efficient genome polishing tool for long-read assembly. Bioinformatics 36, 2253–2255 (2020).

48. N. C. Durand, et al., Juicer Provides a One-Click System for Analyzing Loop-Resolution Hi-C Experiments. Cell Systems 3, 95–98 (2016).

49. O. Dudchenko, et al., De novo assembly of the *Aedes aegypti* genome using Hi-C yields chromosome-length scaffolds. Science 356, 92–95 (2017).

50. M. Xu, et al., TGS-GapCloser: A fast and accurate gap closer for large genomes with low coverage of error-prone long reads. GigaScience 9, giaa094 (2020).

51. H. Cheng, et al., Haplotype-resolved assembly of diploid genomes without parental data. Nat Biotechnol 40, 1332–1335 (2022).

52. S. Ou, et al., Benchmarking transposable element annotation methods for creation of a streamlined, comprehensive pipeline. Genome Biol 20, 275 (2019).

53. M. S. Campbell, C. Holt, B. Moore, M. Yandell, Genome Annotation and Curation Using MAKER and MAKERLP. Current Protocols in Bioinformatics 48 (2014).

54. A. Shumate, S. L. Salzberg, Liftoff: accurate mapping of gene annotations. Bioinformatics 37, 1639–1643 (2021).

55. A. McKenna, et al., The Genome Analysis Toolkit: A MapReduce framework for analyzing next-generation DNA sequencing data. Genome Res. 20, 1297–1303 (2010).

56. C. Camacho, et al., BLAST+: architecture and applications. BMC Bioinformatics 10, 421 (2009).

57. J. Armstrong, et al., Progressive Cactus is a multiple-genome aligner for the thousand-genome era. Nature 587, 246–251 (2020).

58. D. M. Emms, S. Kelly, OrthoFinder: phylogenetic orthology inference for comparative genomics. Genome Biol 20, 238 (2019).

59. G. Hickey, B. Paten, D. Earl, D. Zerbino, D. Haussler, HAL: a hierarchical format for storing and analyzing multiple genome alignments. Bioinformatics 29, 1341–1342 (2013).

60. L. Chen, et al., Large-scale ruminant genome sequencing provides insights into their evolution and distinct traits. Science 364, eaav6202 (2019).

61. G. Talavera, J. Castresana, Improvement of Phylogenies after Removing Divergent and Ambiguously Aligned Blocks from Protein Sequence Alignments. Systematic Biology 56, 564–577 (2007).

62. A. M. Kozlov, D. Darriba, T. Flouri, B. Morel, A. Stamatakis, RAxML-NG: A fast, scalable, and user-friendly tool for maximum likelihood phylogenetic inference. 5.

63. H. Li, Minimap2: pairwise alignment for nucleotide sequences. Bioinformatics 34, 3094–3100 (2018).

64. M. Goel, H. Sun, W.-B. Jiao, K. Schneeberger, SyRI: finding genomic rearrangements and local sequence differences from whole-genome assemblies. Genome Biol 20, 277 (2019).

65. M. Goel, K. Schneeberger, plotsr: visualizing structural similarities and rearrangements between multiple genomes. Bioinformatics 38, 2922–2926 (2022).

66. Y. Wang, et al., MCScanX: a toolkit for detection and evolutionary analysis of gene synteny and collinearity. Nucleic Acids Research 40, e49–e49 (2012).

67. V. Bandi, C. Gutwin, Interactive Exploration of Genomic Conservation. 10.

68. Z. Yang, PAML: a program package for phylogenetic analysis by maximum likelihood. Bioinformatics 13, 555–556 (1997).

69. H. Li, R. Durbin, Inference of human population history from individual whole-genome sequences. Nature 475, 493–496 (2011).

70. K. Nam, et al., Molecular evolution of genes in avian genomes. 17 (2010).

71. J. Terhorst, J. A. Kamm, Y. S. Song, Robust and scalable inference of population history from hundreds of unphased whole genomes. Nat Genet 49, 303–309 (2017).

72. L. Excoffier, et al., *fastsimcoal2*L: demographic inference under complex evolutionary scenarios. Bioinformatics 37, 4882–4885 (2021).

73. N. Patterson, et al., Ancient Admixture in Human History. Genetics 192, 1065– 1093 (2012).

74. M. J. Hubisz, K. S. Pollard, A. Siepel, PHAST and RPHAST: phylogenetic analysis with space/time models. Briefings in Bioinformatics 12, 41–51 (2011).

75. S. Mirarab, et al., ASTRAL: genome-scale coalescent-based species tree estimation. Bioinformatics 30, i541–i548 (2014).

76. E. Sayyari, DiscoVista_ Interpretable visualizations of gene tree discordance. Molecular Phylogenetics and Evolution, 6 (2018).

77. D. Heller, M. Vingron, SVIM: structural variant identification using mapped long reads. Bioinformatics 35, 2907–2915 (2019).

78. K. Wang, M. Li, H. Hakonarson, ANNOVAR: functional annotation of genetic variants from high-throughput sequencing data. Nucleic Acids Research 38, e164– e164 (2010).

79. A. Mayakonda, D.-C. Lin, Y. Assenov, C. Plass, H. P. Koeffler, Maftools: efficient and comprehensive analysis of somatic variants in cancer. Genome Res. 28, 1747–1756 (2018).

80. Z. Zhang, et al., Whole-genome resequencing reveals signatures of selection and timing of duck domestication. GigaScience 7 (2018).

81. R. Liu, et al., Genomic analyses reveal the origin of domestic ducks and identify different genetic underpinnings of wild ducks. 2020.02.03.933069 (2020).

82. T. Zhu, et al., Positive selection of skeleton-related genes during duck domestication revealed by whole genome sequencing. BMC Ecol Evo 21, 165 (2021).

83. H. Li, R. Durbin, Fast and accurate long-read alignment with Burrows–Wheeler transform. Bioinformatics 26, 589–595 (2010).

84. A. R. Quinlan, I. M. Hall, BEDTools: a flexible suite of utilities for comparing genomic features. Bioinformatics 26, 841–842 (2010).

85. C. C. Chang, et al., Second-generation PLINK: rising to the challenge of larger and richer datasets. GigaSci 4, 7 (2015).

86. Fast model-based estimation of ancestry in unrelated individuals (November 24,2022).

87. K. Katoh, D. M. Standley, MAFFT Multiple Sequence Alignment Software Version 7: Improvements in Performance and Usability. Molecular Biology and Evolution 30, 772–780 (2013).

88. B. Q. Minh, et al., IQ-TREE 2: New Models and Efficient Methods for Phylogenetic Inference in the Genomic Era. Molecular Biology and Evolution 37, 1530–1534 (2020).

89. D. Crysnanto, A. S. Leonard, Z.-H. Fang, H. Pausch, Novel functional sequences uncovered through a bovine multiassembly graph. Proc. Natl. Acad. Sci. U.S.A. 118, e2101056118 (2021).

90. W.-W. Liao, et al., A draft human pangenome reference. Nature 617, 312–324 (2023).

91. P. Danecek, et al., Twelve years of SAMtools and BCFtools. GigaScience 10, giab008 (2021).

92. M. Kirsche, et al., “Jasmine: Population-scale structural variant comparison and analysis” (Genomics, 2021) 10.1101/2021.05.27.445886 (December 2, 2022).

93. A. C. English, V. K. Menon, R. Gibbs, G. A. Metcalf, F. J. Sedlazeck, “Truvari: Refined Structural Variant Comparison Preserves Allelic Diversity” (Bioinformatics, 2022) 10.1101/2022.02.21.481353 (December 2, 2022).

94. Z. Wang, et al., Dynamics of transcriptome changes during subcutaneous preadipocyte differentiation in ducks. BMC Genomics 20, 688 (2019).

95. Y. Liao, G. K. Smyth, W. Shi, featureCounts: an efficient general purpose program for assigning sequence reads to genomic features. Bioinformatics 30, 923–930 (2014).

